# Local field potentials primarily reflect inhibitory neuron activity in human and monkey cortex

**DOI:** 10.1101/052282

**Authors:** Bartosz Teleńczuk, Nima Dehghani, Michel Le Van Quyen, Sydney S. Cash, Eric Halgren, Nicholas G. Hatsopoulos, Alain Destexhe

## Abstract

The local field potential (LFP) is generated by large populations of neurons, but unitary contribution of spiking neurons to LFP is not well characterised. We investigated this contribution in multi-electrode array recordings from human and monkey neocortex by examining the spike-triggered LFP average (st-LFP). The resulting st-LFPs were dominated by broad spatio-temporal components due to ongoing activity, synaptic inputs and recurrent connectivity. To reduce the spatial reach of the st-LFP and observe the local field related to a single spike we applied a spatial filter, whose weights were adapted to the covariance of ongoing LFP. The filtered st-LFPs were limited to the perimeter of 800 *μ*m around the neuron, and propagated at axonal speed, which is consistent with their unitary nature. In addition, we discriminated between putative inhibitory and excitatory neurons and found that the inhibitory st-LFP peaked at shorter latencies, consistently with previous findings in hippocampal slices. Thus, in human and monkey neocortex, the LFP reflects primarily inhibitory neuron activity.

## Introduction

The information in neural systems is distributed across a large number of neurons. In order to understand how it is encoded, processed and transformed into actions, we need to monitor activities of a significant fraction of the neuronal population^1^. A popular measure of the population activity is the local field potential (LFP), which represents summed synaptic activity located in small volume around the recording site^2^. Although LFP is easy to record, it has proven notoriously difficult to interpret and model^2,3^. These difficulties partially originate from the complexity of neuronal coding^4,5^, but they also result from the very nature of the LFP signal, which represents the neuronal activity only indirectly through the flow of the extracellular currents^2,6^. This current flow depends on a number of parameters such as the neuronal morphology^7^, synaptic receptors^8^, membrane ion channels^9,10^, electric properties of the tissue^6^, brain area and cortical layer^11^ impeding the interpretation of the resulting LFP signal.

We have begun to understand some of the cellular origins of the LFP signal^3^. In particular, the combined effects of the above factors on the single-neuron contribution to the LFP is a topic of intensive study^2,12^. Each single-neuron spike triggers synaptic input currents in all of its post-synaptic targets, which, along with the corresponding return currents, generates the associated LFP signal also called the unitary LFP. At any time point, the ongoing in-vivo LFP may sum tens of thousands of such unitary signals masking the contributions triggered by a single spike. Unitary LFP has been only characterised in hippocampus, since slice preparations provide the medium for direct current injection and initiation of unitary LFP^13^. These experiments showed unexpectedly strong contribution of interneurons as compared to pyramidal neurons.

The relation of spikes to LFP can be studied in vivo, for example, using spike-triggered LFP average (st-LFP)^14^, which estimates the LFP associated with each spike of a single neuron. Such measures have been used to assess gamma-band synchronisation between neurons^15^, to detect spike locking to phase of oscillatory LFP^16^, to characterise the synaptic connectivity^17^ and to study travelling cortical waves^18^. In addition, the st-LFP is modulated by the waking state^16^ and stimulus contrast^19^. However, the unitary LFP could not be identified using this technique, because st-LFP can not discern it from the ongoing LFP activity and recurrent activity in the network^2,20^. Therefore, we introduce a spatial filtering technique that helps to separate the effects of a single neuron from the non-specific LFP components common to the local population.

Using this technique we aimed to differentiate the contributions of interneurons and pyramidal neurons to LFP recorded from humans and monkeys. We anticipated that the long-duration recordings with dense grid of electrodes (Utah array)^21,22^ would allow us to separate unitary LFP of putative interneurons and pyramidal neurons. Since neuronal morphology and connectivity affect the LFP^7,11^, we expected that these two types of neurons should be associated with diverse LFP contributions. We found that both inhibitory and excitatory neurons are associated with st-LFPs dominated by spatially and temporally broad components. We then estimated spatial filters adapted to the structure of ongoing LFP, which allowed us to focus on the focal contributions instead. Using these methods we could demonstrate for the first time that the post-synaptic currents initiated by inhibitory interneuron spikes are the dominant generators of focal LFP.

## Results

We investigated the local field potential (LFP) contribution associated with a single spike in human and monkey cortex. The data were recorded from the temporal cortex of patients who underwent a surgical procedure for the localisation of the epileptic foci^21^ and from the dorsal premotor cortex (PMd) of macaque monkey^5^ (Figure 1A). The LFP and spiking activity (Figure 1B) were recorded with a 10-by-10 array of intracortical electrodes (Utah array, interelectrode distance 400 *μ*m). Spikes of single neurons were sorted by semi-automatic clustering (see Methods). In total, we analysed data from 206 neurons in two human subject and 152 in one monkey (the detailed information on the number of neurons and spikes is available in Supplementary Table 1). The relation between spikes and LFP was estimated with the spike-triggered LFP average (st-LFP), that is the average of short LFP segments centered around each spike time. This procedure was applied to LFP signals from all electrodes using spikes of a given neuron as the trigger. Thus we obtained a spatio-temporal map of the LFP components coincident with a spike of a given neuron (Figure 1C).

**Figure 1:**
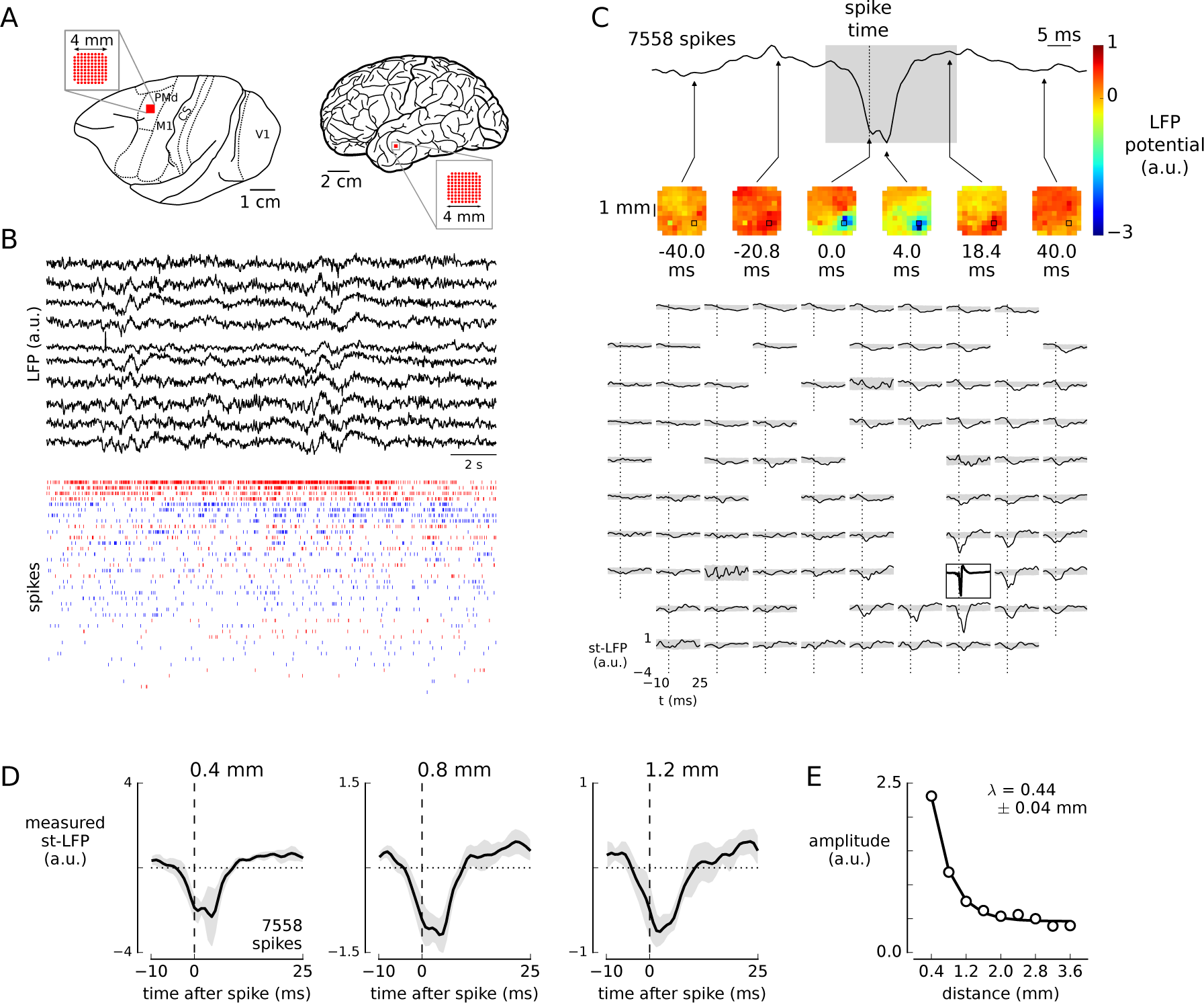
Spikes of single neurons are associated with spatially diffuse and non-causal LFP patterns. (**A**) Local field potentials and spikes were measured in the premotor cortex of a macaque monkey (top) and temporal cortex of human subjects (bottom) using the Utah arrays. (**B**) LFP (top, subset of LFPs recorded simultaneously from macaque premotor cortex) and spikes (bottom, subset of neurons) obtained from Utah array. Neurons were classified into regular spiking (bottom, blue) and fast spiking (red) types based on spike waveshape. (**C**) Spatio-temporal spike-triggered LFP average (st-LFP) in human temporal cortex. Top: Average of the st-LFPs (band-pass filtered 15 – 300 Hz, average of 7558 spike-triggered segments) from the electrodes neighbouring with the trigger neuron. Middle: A color map of st-LFP amplitudes at selected time lags around the spike. The values for missing electrodes were replaced with the average of the neighbouring electrodes. Bottom: The st-LFP from all valid electrodes of the array plotted in time (plotting window adjusted to the gray-shaded segment in top panel). The st-LFP at the neuron position (black rectangle) was replaced with the spike waveform (amplitude normalised). Most st-LFPs express non-causal components preceding the spike (spike onset shown with vertical dotted line). The gray-shaded area represents 95% confidence intervals calculated from jittered spikes (1000 repetitions, gaussian jitter 100 ms). (**D**) st-LFPs triggered on spikes of a single neuron (same as shown in (C)) and averaged for all electrodes separated by the same distance from the neuron (3 distances: 0.4 mm, 0.8 mm, 1.2 mm shown from left to right). The 95% confidence intervals (gray-shaded area) were calculated as ±1.96×s.e.m. (**E**) The st-LFP trough amplitude as a function of the distance from the neuron to the LFP electrode. The data points were fitted with an exponential *A*exp(-*x*/*λ*) + *C*, where *x* is the distance and *λ* is the space constant (fitted value ± SD in the top-right corner).

The estimated st-LFPs are not confined to local neighbourhood of the trigger neuron, but they spread rapidly through the entire array (Figure 1C, middle and bottom): the LFP deflection related to the spike reaches most electrodes within 4 ms after the spike onset (see the heatmap in Figure 1C). To measure the spatial extent, we averaged st-LFPs from electrodes with the same distance from the trigger neuron (Figure 1D). We found that these spike-related LFP components could persist over distances larger than 1 mm away from the neuron that initiated them (Figure 1D, gray-shaded area denotes the confidence intervals). The amplitude of the negative peak (trough) decayed exponentially from the trigger neuron with the space constant of *λ* = 0.44±0.04 mm (mean ± standard deviation, Figure 1E). In addition, many components, even at distant electrodes, appear nearly simultaneously with a spike of the reference neuron or may even precede the spike (as shown by the fact that confidence intervals prior to spike onset do not cross the zero line, Figure 1D). Such non-local and non-causal components can not be interpreted as the field generated directly by the active neuron (via the post-synaptic potentials), because most neurons synapse at distances no more than 1 mm^23,24^.

Similar results were obtained from other neurons in both human and monkey subjects. To compare results across recordings we averaged the st-LFP across equi-distant electrodes and across neurons (see Methods). We grouped the neurons into two populations: putative interneurons (fast spiking, FS) and pyramidal neurons (regular spiking, RS), which were discriminated on the basis of spike waveshape^21,25^ (Figure 2). The rate of spatial decay varied slightly across the subjects: from 0.48 mm to 0.71 mm for RS and from 0.22 mm to 0.57 mm for FS neurons. For two subjects (one human subject and macaque) the space constants for RS-and FS-based st-LFPs differed significantly (t-test, p<0.01), but the direction of the difference was not consistent across subjects (the space constant, *λ*, was larger for RS neurons in the second human subject, whereas in monkey the relation was inverted). We also compared the absolute values of the st-LFP amplitudes at the distances 0.4 – 2 mm, but we did not find significant differences in any of the subjects (except the second night of the first human subject, Supplementary Figure 7A; bootstrap test, p < 0.05).

**Figure 2:**
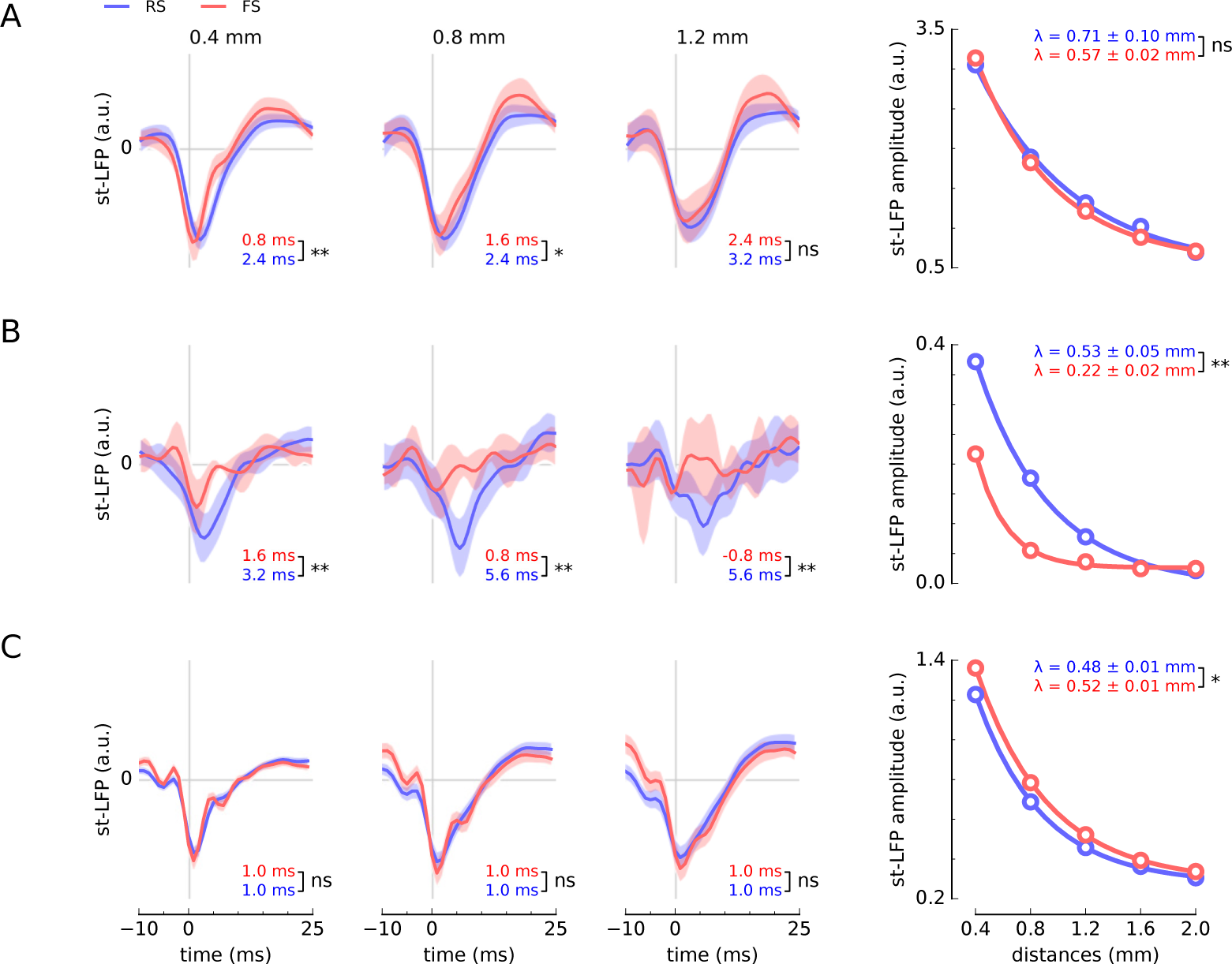
Spatial and temporal st-LFP components across neurons and subject. (**A** and **B**) Human subjects. (**C**) Monkey. *Three left-most panels*: st-LFPs triggered by spikes of regular spiking (RS, blue line) or fast spiking (FS, red line) neurons at 0.4, 0.8 and 1.2 mm from the trigger neuron. The st-LFPs were averaged both over neurons and electrodes. The trough latencies of RS and FS neurons (values given in bottom-right corner) are significantly different for short distances in one human subject and monkey. The shaded areas represent the 95% confidence intervals of the respective st-LFPs (±1.96×s.e.m.). *Right-most panel*: The decay of trough amplitudes with distance. The RS/FS space constants *λ* (coefficient SD given in top-right corner) as determined by fitting an exponential function (solid line) to the estimated amplitudes (circles) were significantly different (tested using t-test) for second human subject (B) and monkey (C). n.s: not significant, *: *p* < 0.05, **: *p* < 0.01, * * *: *p* < 0.001.

To further elucidate the relation of RS and FS neurons to the LFP, we also determined the latencies of the deepest trough in the estimated st-LFP (Figure 2, bottom-right corners of the panels). We found that at 0.4 mm from the trigger neuron the st-LFPs reached the minimum at 0.8 – 1.6 ms (FS neurons) or 1.0 – 3.2 ms (RS neurons) after spike onset. The trough latencies of RS and FS neurons were significantly different for both human subjects (bootstrap test, p < 0.01), so we can conclude that the FS neurons correlated with LFP potentials at shorter latencies compared to RS neurons. In monkey premotor cortex the latency for FS and RS neurons did not differ significantly.

We hypothesised that the spatial spread of the st-LFP results partially from the broadening of the unitary LFP either by the passive propagation of the electric field (“volume conduction”) or other neurons firing spikes at correlated times. To recover the focal contribution of the trigger neuron we applied spatial filters that decorrelated (“whitened”) ongoing LFP signals in space^26^ keeping the spike-related components. We validated the method on a simple linear model representing the LFP signal as a sum of post-synaptic contributions from a neuronal population (see Supplementary Methods). We demonstrate that the obtained st-LFP is broader spatially than the post-synaptic kernel (unitary LFP). However, the whitening procedure recovers the unitary LFP by reducing spurious spatial correlations (Supplementary Figure 1).

The filters estimated from the experimental data are similar to the second spatial derivative (laplacian), but they also feature additional off-center components (Figure 3A). When applied to the st-LFP they suppressed the global components present in the entire array and kept only small components localised around the neuron (Figure 3B, right, and Supplementary Figure 5). The whitened st-LFPs (wst-LFPs) reached no further than 1 mm from the trigger neuron so they may originate from the synaptic currents directly evoked by its spikes (Figure 3D, right). Similarly, focal LFP contributions were recovered in the second human subject and macaque dorsal premotor cortex (Figure 3E-F). The space constants of wst-LFP amplitude decay were half as long as the non-whitened st-LFP: for RS neurons they ranged from 0.20 to 0.25 mm and for FS neurons from 0.15 mm to 0.20 mm. However, the difference between space constants of wst-LFP of RS and FS neurons were not found to be significantly different (t-test, p>0.05). Additionally, we compared the wst-LFP obtained in the awake periods with slow-wave periods of human subject 1; we did not find significant differences in the amplitudes and time courses of these traces, but we identified significant state-related differences in space constants for RS neurons (Supplementary Figure 4).

**Figure 3:**
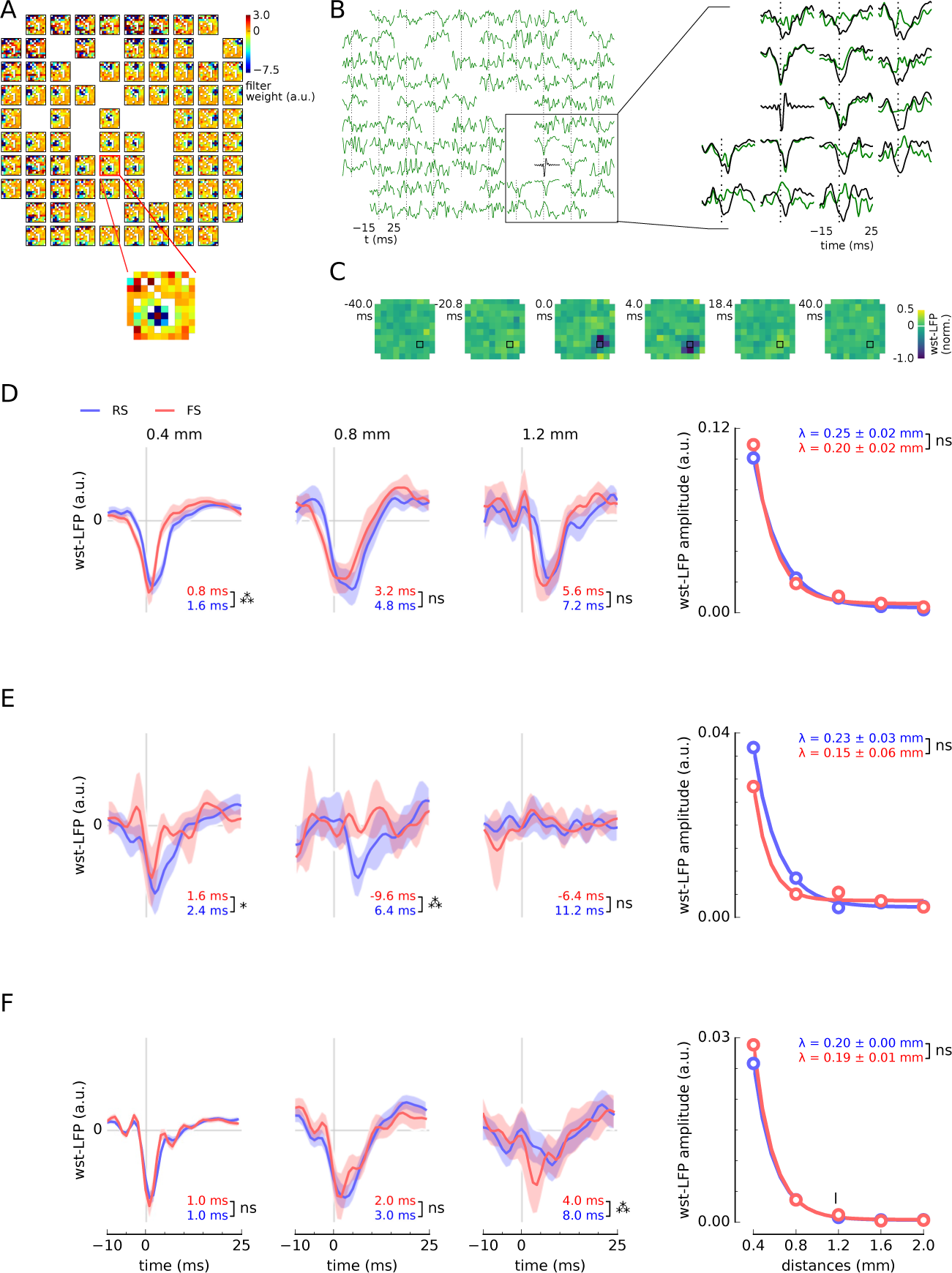
The focal st-LFP of fast spiking (FS) and regular spiking (RS) neurons recovered by spatial decorrelation (whitening). (**A**) Spatial filters designed to decorrelate (whiten) the LFP signals. Colors show filter weights associated with each electrode. Inset : Scaled-up heatmap of weights to whiten a single st-LFP (single row of whitening matrix). (**B**) Whitened st-LFPs (wst-LFPs) of a single neuron (green) compared with the non-whitend st-LFP (black). Right : close-up of the whitened st-LFPs enclosed in black rectangle. Most st-LFPs are suppressed after spatial whitening and only st-LFPs directly adjoining the neuron are conserved. (**C**) Spatial maps of whitened st-LFP (compare with Figure 1C). (D-F) The population-averaged st-LFPs after whitening for human subject 1 (**D**), human subject 2 (**E**) and monkey (**F**). Three panels from left : Whitened st-LFPs averaged across neurons and electrodes at three distances from the trigger neuron (0.4 mm, 0.8 mm, 1.2 mm; for details see legend of Figure 2). Right-most panel : st-LFP trough amplitude as a function of the distance between the neuron and the LFP electrode (cf. Figure 1E). The star indicates significant differences in the trough amplitude between RS and FS neurons at the respective distance. n.s: not significant, *: *p* < 0.05, **: *p* < 0.01, * * *: *p* < 0.001.

**Figure 4:**
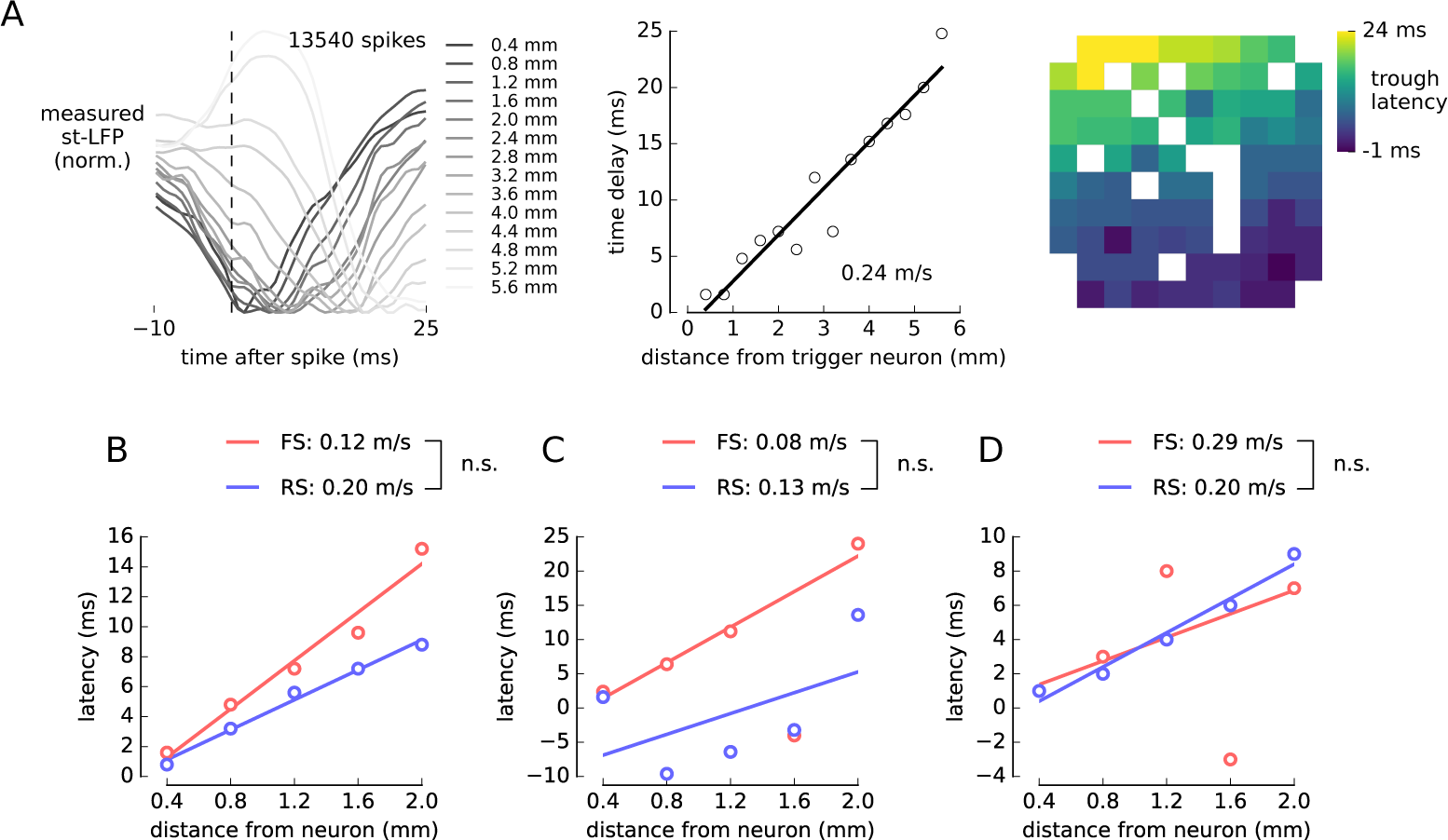
Propagation of st-LFP in human and monkey. (**A**) Propagation of the non-whitened st-LFP across the electrode array. From left to right : single-neuron st-LFPs averaged over all electrodes with the same distance from the trigger neuron; the latency of averaged st-LFP troughs (dots) and the linear fit (solid line); the latency of st-LFPs from each electrode as a heatmap (blank squares are due to missing electrodes). The latencies increase with the distance supporting the hypothesis of spike-evoked LFP propagation. (**B**) The propagation of whitened and population-averaged st-LFPs in human. Dependency of latency on distance, separately for each neuron type, is plotted against the distance from the trigger neuron (circles) and fitted with a linear function (solid line). The speeds of st-LFPs propagation for RS and FS neurons were calculated from the inverse of the slope (number at the top). The speed differences tested using bootstrap method could not be shown significantly different (bracket next to speed values). (**C**) Second human subject. (**D**) Monkey. The wst-LFP traces for which the latencies were calculated are shown in Figure 3 and Supplementary Figure 3. n.s: not significant, *: *p* < 0.05, **: *p* < 0.01, * * *: *p* < 0.001.

We also found that in their close neighbourhood both the RS and FS neurons produced negative-going wst-LFPs of similar amplitudes (Figure 3D-F, right). In both human subjects and at 0.4 mm from the trigger neurons, wst-LFP of FS population peaked consistently before the RS-based wst-LFPs (Figure 3D-F). In monkey recordings the latency difference was only significant at 1.2 mm from the trigger neuron (Figure 3F). The latency differences could be explained by a di-synaptic contribution of the RS neurons: RS neurons would excite the inhibitory interneurons, which in turn would contribute to the LFP. In this model the shift in st-LFP latency reflects the delay due to synaptic transmission and axonal delays. Importantly, this explanation also accounts for the fact that the excitatory and inhibitory wst-LFPs have the same polarity.

The latency of the (non-whitened) st-LFPs increased gradually with the distance from the trigger neuron (Figure 4A, left). This is shown more precisely when plotting the latency of the trough against the electrode distance (Figure 4A, middle). The linear dependency between these measures suggests propagation with constant speed, which can be estimated from the inverse slope of the linear fit. The dispersion of some points from the best-fitting line can be attributed to the error in the estimation of the latency. For the st-LFPs averaged over all neurons of each type (Supplementary Figure 6) the estimated propagation speed was 0.36 m/s for both FS and RS neurons. For the second human subject and monkey it was not possible to determine the propagation speed, because the latency either did not depend linearly on distance, or the dependence was inverted (decreasing latency with distance, Supplementary Figure 6).

These distortions of the latency values could arise due to electric phenomena occurring in the extracellular medium. In particular, the estimated propagation speeds might reflect the synaptic propagation of action potentials (axonal conduction times and synaptic delays), but passive phenomena such as low-pass filtering by the medium^27^ or by dendritic structure^7^ could also affect the st-LFP. To focus on the synaptic phenomena, we analysed the propagation in the whitened st-LFP. We found that in one human subject the propagation was slower in the wst-LFP (FS: 0.12 m/s, RS: 0.20 m/s) in comparison to the non-whitened st-LFP (Figure 4B). This may be due to the fact that the spatial filtering removes the effects of currents passively conducted through the tissue (volume conduction), which can propagate much faster than the synaptically-transmitted signals^27^. The combination of the nearly instantaneous (volume-conducted), and delayed (synaptically-propagated) st-LFP components may explain the higher overall propagation velocity in the non-whitened st-LFP. To check whether this propagation speed does not vary across time, we repeated the same analysis for the second day of the recording and we obtained similar estimates (Supplementary Figure 7, FS: 20 m/s, RS: 19 m/s).

Owing to large dispersion of the latency measurements (Supplementary Figure 6), it was not possible to estimate the propagation speed of non-whitened st-LFP for the second human subject and the monkey. However, the estimated latencies where more consistent after the application of whitening filters, so that the propagation speed could be also determined for the FS neurons in the second human subject (0.08 m/s) and both RS (0.20 m/s) and FS (0.29 m/s) neurons in monkey. Interestingly, propagation speeds in all three subjects range between 0.08 and 0.29 m/s consistently with the action potential propagation speed in unmyelinated fibers^28^.

## Discussion

We studied the relationship between neuronal activity and its surrounding electrical field by means of simultaneous recordings of single-unit spikes and the LFP signal. We found that the spike-triggered LFP can be described by a focal component, which reflects single-neuron activity, and a diffuse component related to the baseline LFP correlations. Data-driven spatial filters recovered the focal LFP around each neuron, which spreads at distances consistent with cortical anatomy and connectivity. The peak latencies and propagation of such whitened st-LFP depended on the type of neuron (inhibitory vs excitatory) used as the trigger. Specifically, the inhibitory neurons provide the largest contribution to the LFP in their close neighbourhood (< 1 mm).

The spike-triggered LFP is an estimate of correlation between the (continuous) LFP signal and the (point-like) spike trains. As a correlation measure it can not differentiate the causal contributions of the spike and its post-synaptic consequences from the incidental correlations between LFP and spikes. In line with the argument, we identified st-LFP components that were both non-causal (i.e. preceded the spike onset) and non-local (appearing simultaneously in distant electrodes, Figure 1C). These components are not specific to the activity of the “trigger” neuron, but characterise the local population of neurons and electric properties of neural tissue^2^.

Notwithstanding, we recovered the local correlates of spikes in the LFP using spatial (whitening) filters adapted to the covariance structure of ongoing LFPs (Figure 3A-C). Similar techniques have been used as a pre-processing step of blind source separation methods^26^ and to study neuronal responses to natural stimuli^29^.

Importantly, we found that the whitened st-LFP (wst-LFP) were much sharper spatially decaying within 1 mm from the trigger neuron. This spatial width corresponds with the reach of axon branches for the locally connected neurons (interneurons). In addition, the temporal profile of the wst-LFP resembles the trace of post-synaptic currents mediated by GABA and AMPA receptors. Although st-LFP advances in space at speed only attributable to the passive electric field propagation, the latency of the wst-LFP is consistent with the delays imposed by synaptic transmission and the propagation of the action potential along an unmyelinated axon. All of these lines of evidence converge on the idea that the wst-LFP represents the electric field due to post-synaptic currents initiated by spikes of the trigger neuron, that is the unitary LFP. We found that the latencies, amplitude decays, and propagation speeds of wst-LFP are consistent across subjects, cortical areas and species (humans and macaque monkeys). This suggests that the described aspects of the LFP generation by the post-synaptic potentials of inhibitory and excitatory neurons are a general feature of the cortical tissue.

In spite of being largely consistent with the expected features of unitary LFP, the st-LFP also shows some small components preceding the spike, which could not be interpreted as causal synaptic consequence of the spike on its post-synaptic targets. However, the methods used to estimate the st-LFP might introduce several artifacts: 1) The smearing of the unitary contribution due to the time-domain filtering implemented both in hardware and in offline analysis; 2) Contribution from synaptic inputs that produced the spike; 3) Imprecise estimation and inversion of the covariance matrix. An alternative, and possibly more direct, method of assessing the spike contribution to LFP would consist in triggering a spike externally at random times (for example, by means of direct current injection) and recording the associated LFP signal. This technique might dissociate the times of spiking from the neurons’s synaptic inputs and activity of other neurons. However, such recordings in intact tissue are technically challenging and to the best of our knowledge have not been performed in monkey and humans.

Most of our results support the hypothesis that st-LFP represents the unitary LFP. However, due to the correlative nature of the whitened and non-whitened st-LFP, we can not exclude alternative interpretations. For example, the axon-propagating action potential could produce similar LFP contribution, albeit at shorter spatial ranges constrained by the axonal arborisation^30^. It has been also suggested that the LFP is at least partially generated by dendritic spikes^31^, which could constitute another source of the observed st-LFP traces. It remains to be investigated whether and to what degree these phenomena contribute to the st-LFP obtained in this study.

We also note that the recordings in humans were done invasively in epileptic cortex. Although the Utah array was not placed in the epileptic focus, some pathological activity was still present. Therefore, for the analysis we only chose the data from non-epileptic periods. The agreement between the healthy tissue of macaque motor cortex and the epileptic human data further confirms the applicability of these results to normal physiological conditions.

The discrimination of neuron types used in this study is based on the assumption that inhibitory interneurons produce thin spikes. Previously, we have shown that in human recordings, the separation provided by spike shape was mirrored by excitatory or inhibitory functional interactions identified from short-latency cross-correlograms. Thus spike-waveform features enable us to classify neurons as inhibitory and excitatory^21^. We have followed a similar procedure for the monkey recordings^32^. This procedure is well established in rodents^33,34^, but has some limitations in recordings from cats^35^ and primates^36,37^. In particular, rarely some excitatory cells of motor cortex may exhibit narrow spikes^36^. In addition, some subtypes of inhibitory, non-fast-spiking interneurons show broad waveforms^38^. However, as these cells represent at most half of the GABAergic neuronal population^34^ and that, overall, inhibitory interneurons represent about 20% of all cortical neurons, the false positive rate of excitatory cell classification is at most 10%. Such misclassifications could affect quantitatively some of our results, but they will not change our conclusions. In our dataset the inhibitory interneurons seem to be overrepresented compared to the 20% proportion of cortical neurons (22% of inhibitory neurons in all human subjects and 35% in monkey), which could reflect the sampling bias in our recordings.

Our results are consistent with the finding that the main contribution to the LFP in hippocampal slices (the unitary field potentials) is due to inhibitory neurons, while the contribution from excitatory neurons is mediated by interneurons, di-synaptically^13^. Two of our findings suggest that these conclusions are valid for human and monkey: the polarities of the wst-LFPs of putative inhibitory and excitatory neurons are the same; and the excitatory contribution to the LFP lags behind the inhibitory (Figure 3D-E). Since the inhibitory and excitatory post-synaptic currents have opposite directions, they should produce LFP of inverse polarities. The fact that the wst-LFPs have the same polarities suggests that excitatory unitary LFP are actually produced di-synaptically by mediating interneurons. This conclusion is further supported by the larger amplitude of inhibitory wst-LFPs that was obtained in one subject (second night of human subject 1, Supplementary Figure 7B).

Alternatively, same polarities of wst-LFP could be also obtained if the different neuron types make synapses in different layers. However, this hypothesis would not account for the shorter latency of the wst-LFPs produced by the inhibitory neurons. The extra delay observed in the putative excitatory wst-LFPs might reflect the synaptic delay required for the di-synaptic activation of mediating interneurons. Similar results were obtained in the macaque monkey, but we found significant differences in the latency only at 1.2 mm from the trigger neuron.

Similar mechanism of LFP generation by inhibitory and excitatory neurons was suggested by Bazelot et al.^13^, who argued that the differences between their unitary LFPs are related to the axonal arboristation and distribution of synaptic terminals. The results of the present study are consistent with this interpretation, but our experimental protocol differs in terms of the preparation used (in vivo vs in vitro experiments) and recorded brain area (premotor/temporal cortex vs hippocampus). Another source of uncertainty is the distribution of axons and synapses across cortical layers: Ideally one should have estimates of the spatial distribution of synapses. Unfortunately such data are not yet available for human cortex and we can not directly address the origin of the neuron-type-related differences in the st-LFP.

The spike-triggered LFP remains an essential method in answering how activities of single neurons are embedded in ongoing rhythms^19^. Here, we demonstrated that st-LFP can be used to assess the specific contribution of the single neurons to the LFP and, indirectly, their synaptic connectivity. Our results suggest also that such contribution might be conserved across brain areas and states and that it is stable over timescales of several hours. Future work might clarify whether it might undergo dynamic changes, for example, during learning. The approach that we adapted here provides a new way to investigate the biophysical link between microscopic and macroscopic scales of cortical organisation.

## Methods

### Experimental methods

Human recordings were acquired from inpatient invasive monitoring of two female patients (52 and 24-year-old) with focal temporal lobe epilepsy. Two nights of recording (each was 12-hour-long) were collected and analysed for the first patient. Most results are reported for the first night only (Figures 2A, 3D and 4B); the second night was used for the confirmation of the stability of the results across time (Supplementary Figure 7). A third patient recorded from the same study was not analysed here due to large fraction of seizure-like activity. The neuroprobe (Utah array) was placed in layer II/III of the middle temporal gyrus. This array is silicon-based, made up of 10x10 (96 recording) microelectrodes with 400 *μ*m spacing, covering an area of 4 × 4 mm. Data were sampled at 30 kHz (Cerebrus Blackrock Microsystems). The LFP signals were obtained from the raw recordings by low-pass filtering and subsampling to 1250 Hz. The epileptic-like activity (interictal spikes and seizure periods) was detected by visual inspection of the LFP and EEG signals and rejected from the analysis. Single units were detected in the 30-kHz recordings and discriminated using standard clustering methods (KulstaKwik, http://klustakwik.sourceforge.net) and manually processed using Klusters software. The data were recorded under normal behaviour (no specific task was administered during this recording) during a single 12-hour-long session covering both awake and sleep periods (as seen in EEG and video recordings). The sleep and awake periods were combined to obtain better statistics. The full experimental protocol can be found in Peyrache et al.^21^.

The monkey recordings were performed during the night in the premotor cortex of macaque monkey (Macaca mulatta) implanted with Utah arrays described above. During a recording session, signals from 96 electrodes were amplified (gain of 5,000), band-pass filtered between 0.3 Hz and 7.5 kHz, and recorded digitally (14-bit) at 30 kHz per channel using a Cerebus acquisition system (Blackrock Microsystems). Spike-waveform data were sorted offline (Plexon, Dallas, TX) using a user-defined template. All spike waveforms whose mean squared error from this template fell below a user-defined threshold were classified as belonging to that unit. The full experimental protocol can be found in Dickey et al.^22^.

We discriminated the putative regular-spiking (RS) and fast-spiking (FS) neurons based on their extracellular spike features. In brief, the averaged spike waveforms were described by 4 features (peak-to-valley amplitude, positive and negative half-width, positive-to-negative interval), which were then used for clustering (K-means) analysis. In humans, only neurons that produced welldefined clusters (quantified by the distance from the decision boundary) were used in further analysis. In addition, for some of neurons the type was confirmed by means of the interaction type identified using pairwise correlograms. In monkey, the stability of the neuron type identification was quantified during three separate recording sessions – only neurons that were consistently classified as either inhibitory or excitatory were used for further analysis. The details of the procedure for human and monkey data were described elsewhere^21,25^.

Approval for all human experiments involving recordings of single unit activity in patients was granted by the Institutional Review Boards of Massachusetts General Hospital/Brigham & Women’s Hospital in accordance with the Declaration of Helsinki and required informed consent was obtained from each participant. For the primate experiments, all of the surgical and behavioral procedures were approved by the University of Chicago’s IACUC and conform to the principles outlined in the Guide for the Care and Use of Laboratory Animals (NIH publication no. 86–23, revised 1985; IACUC Approval number: 71565).

### Data analysis

We calculated the spike-triggered average of the LFP (st-LFP) in order to estimate the contribution of a single spike to LFP^29^. First, we calculated the average of short segments of the LFP centered around each spike time of a single trigger neuron. Next, we repeated this process across LFP electrodes to obtain a spatio-temporal st-LFP:

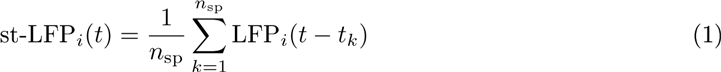

where *i* = 1.96 are the indices of the electrodes and 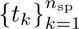 are all *n*_sp_ spike times of the trigger neuron. To avoid spike artifacts, the LFP signal from the electrode that recorded the spikes of the trigger neuron was removed from the data (see Supplementary Figure 2). For the purpose of visualisation (Figure 1C and Figure 3C) the missing electrodes were replaced with the average of neighbour electrodes.

We averaged the single st-LFPs across neurons and electrodes. First, for each neuron we averaged the st-LFPs from all electrodes that were separated from the trigger neuron by the same distance (Mahalanobis distance). We repeated this procedure for all distances from 0.4 to 3.2 mm. Secondly, for each distance we averaged the electrode-averaged st-LFPs across neurons obtaining the population-averaged st-LFP. Note that by averaging st-LFPs at constant distances we assumed that the st-LFP is isotropic neglecting the possible direction specificity. This assumption might not be valid for single-neuron st-LFPs (see Figures 1C, 3B-C, 4A), but may be justified in the population average. We estimated the trough amplitudes and latencies in the population-averaged st-LFP (Figures 4B-D) by finding the global minimum of the st-LFP in the time-window of [-10, 15] ms.

To whiten the st-LFPs we first calculated the covariance matrix of the ongoing band-pass filtered (15 – 300 Hz) LFP signals (**C**_LFP_)_*ij*_ = 〈LFP_*i*_(*t*)LFP_*j*_(*t*)〉_*t*_ (where 〈·〉_*t*_ denotes temporal averaging). The whitening matrix **W** is calculated by the inverse square root of **C**_LFP_^26^:

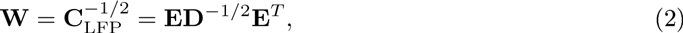

where **E** is a matrix of eigenvectors of **C**_LFP_ and **D** is a diagonal matrix with inverse square roots of eigenvalues *λ*_*i*_ on its diagonal 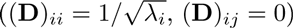.

Given the matrix **W** the whitening operation amounts to the matrix product of **W** with the spike-triggered LFP:

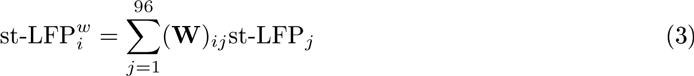

where st-LFP^*ω*^s are the whitened st-LFPs.

In pre-processing steps, the LFP signals were band-pass filtered 15 – 300 Hz. The filtering was applied in Fourier domain where filter response was 1 for all frequencies in the pass-band and 0 in stop-band. To avoid ringing artifacts, at the corner frequencies the response decayed to 0 with a profile of a Gaussian with width of 10 Hz.

## Acknowledgments

Research funded by Centre National de la Recherche Scientifique (CNRS, France), European Community Future and Emerging Technologies program (BrainScales FP7-269921; The Human Brain Project FP7-604102), National Institutes of Health (NIH grants 5R01NS062092, R01EB009282 and R01NS045853, R01MH099645) and Office of Naval Research (MURI grant number N00014-13-1-0672).

## Author contributions

B.T and A.D conceived the study. S.C, E.H and N.H conducted the experiments. B.T analyzed the experimental data with contributions from N.D and M.LVQ. B.T, N.D and A.D interpreted the results and wrote the paper with contributions from all authors.

## References

[1] Georgopoulos, A. P., Schwartz, A. B. & Kettner, R. E. Neuronal population coding of movement direction. Science 233, 1416–1419 (1986).

[2] Einevoll, G. T., Kayser, C., Logothetis, N. K. & Panzeri, S. Modelling and analysis of local field potentials for studying the function of cortical circuits. Nat Rev Neurosci 14, 770–785 (2013).

[3] Buzsáki, G., Anastassiou, C. A. & Koch, C. The origin of extracellular fields and currents–EEG, ECoG, LFP and spikes. Nat. Rev. Neurosci. 13, 407–420 (2012).

[4] Denker, M. et al. The Local Field Potential Reflects Surplus Spike Synchrony. Cereb. Cortex 21, 2681–2695 (2011).

[5] Saleh, M., Takahashi, K. & Hatsopoulos, N. G. Encoding of coordinated reach and grasp trajectories in primary motor cortex. J. Neurosci. 32, 1220–1232 (2012).

[6] Bédard, C., Kröger, H. & Destexhe, A. Modeling Extracellular Field Potentials and the Frequency-Filtering Properties of Extracellular Space. Biophys J 86, 1829–1842 (2004).

[7] Lindén, H., Pettersen, K. H. & Einevoll, G. T. Intrinsic dendritic filtering gives low-pass power spectra of local field potentials. J Comput Neurosci 29, 423–444 (2010).

[8] Mazzoni, A. et al. Computing the local field potential (LFP) from integrate-and-fire network models. PLOS Comput Biol 11, e1004584 (2015).

[9] Reimann, M. W. et al. A biophysically detailed model of neocortical local field potentials predicts the critical role of active membrane currents. Neuron 79, 375–390 (2013).

[10] Ness, T. V., Remme, M. W. & Einevoll, G. T. Active subthreshold dendritic conductances shape the local field potential. J. Physiol. (Lond.) (2016).

[11] Lindén, H. et al. Modeling the spatial reach of the LFP. Neuron 72, 859–872 (2011).

[12] Hagen, E. et al. Hybrid scheme for modeling local field potentials from point-neuron networks. arXiv 1511.01681 (2015).

[13] Bazelot, M., Dinocourt, C., Cohen, I. & Miles, R. Unitary inhibitory field potentials in the CA3 region of rat hippocampus. J. Physiol. (Lond.) 588, 2077–2090 (2010).

[14] Teleńczuk, B., Baker, S. N., Kempter, R. & Curio, G. Correlates of a single cortical action potential in the epidural EEG. Neuroimage 109, 357–367 (2015).

[15] Gray, C. M. & Singer, W. Stimulus-specific neuronal oscillations in orientation columns of cat visual cortex. Proc. Natl. Acad. Sci. U.S.A 86, 1698–1702 (1989).

[16] Destexhe, A., Contreras, D. & Steriade, M. Spatiotemporal analysis of local field potentials and unit discharges in cat cerebral cortex during natural wake and sleep states. J. Neurosci. 19, 4595–4608 (1999).

[17] Swadlow, H. A., Gusev, A. G. & Bezdudnaya, T. Activation of a cortical column by a thalamocortical impulse. J. Neurosci. 22, 7766–7773 (2002).

[18] Nauhaus, I., Busse, L., Carandini, M. & Ringach, D. L. Stimulus contrast modulates functional connectivity in visual cortex. Nat. Neurosci. 12, 70–76 (2009).

[19] Ray, S. & Maunsell, J. H. R. Network rhythms influence the relationship between spike-triggered local field potential and functional connectivity. J. Neurosci. 31, 12674–12682 (2011).

[20] Nauhaus, I., Busse, L., Ringach, D. L. & Carandini, M. Robustness of traveling waves in ongoing activity of visual cortex. J. Neurosci. 32, 3088–3094 (2012).

[21] Peyrache, A. et al. Spatiotemporal dynamics of neocortical excitation and inhibition during human sleep. Proc. Natl. Acad. Sci. U.S.A. 109, 1731–1736 (2012).

[22] Dickey, A. S., Suminski, A., Amit, Y. & Hatsopoulos, N. G. Single-unit stability using chronically implanted multielectrode Arrays. J. Neurophysiol. 102, 1331–1339 (2009).

[23] Holmgren, C., Harkany, T., Svennenfors, B. & Zilberter, Y. Pyramidal cell communication within local networks in layer 2/3 of rat neocortex. J. Physiol. (Lond.) 551, 139–153 (2003).

[24] Levy, R. B. & Reyes, A. D. Spatial profile of excitatory and inhibitory synaptic connectivity in mouse primary auditory cortex. J. Neurosci. 32, 5609–5619 (2012).

[25] Dehghani, N. et al. Dynamic Balance of Excitation and Inhibition in Human and Monkey Neocortex. Sci Rep 6, 23176 (2016).

[26] Hyvärinen, A., Karhunen, J. & Oja, E. Independent Component Analysis (John Wiley & Sons, New York, 2004).

[27] Bédard, C. & Destexhe, A. Modeling local field potentials and their interaction with the extracellular medium. In Brette, R. & Destexhe, A. (eds.) Handbook of Neural Activity Measurement, 136–191 (Cambridge University Press, Cambridge, 2012).

[28] Debanne, D., Campanac, E., Bialowas, A., Carlier, E. & Alcaraz, G. Axon Physiology. Physiol Rev 91, 555–602 (2011).

[29] Schwartz, O., Pillow, J. W., Rust, N. C. & Simoncelli, E. P. Spike-triggered neural characterization. J Vision 6, 484–507 (2006).

[30] Mitzdorf, U. Current source-density method and application in cat cerebral cortex: investigation of evoked potentials and EEG phenomena. Physiol Rev 65, 37–100 (1985).

[31] Nicholson, C. & Llinas, R. Field potentials in the alligator cerebellum and theory of their relationship to Purkinje cell dendritic spikes. J. Neurophysiol. 34, 509–531 (1971).

[32] Quyen, M. L. V. et al. High-frequency oscillations in human and monkey neocortex during the wake–sleep cycle. PNAS 113, 9363–9368 (2016).

[33] Kawaguchi, Y. Groupings of nonpyramidal and pyramidal cells with specific physiological and morphological characteristics in rat frontal cortex. J. Neurophysiol. 69, 416–431 (1993).

[34] Jiang, X. et al. Principles of connectivity among morphologically defined cell types in adult neocortex. Science 350, aac9462 (2015).

[35] Nowak, L. G., Azouz, R., Sanchez-Vives, M. V., Gray, C. M. & McCormick, D. A. Electrophysiological Classes of Cat Primary Visual Cortical Neurons In Vivo as Revealed by Quantitative Analyses. J Neurophysiol 89, 1541–1566 (2003).

[36] Vigneswaran, G., Kraskov, A. & Lemon, R. N. Large Identified Pyramidal Cells in Macaque Motor and Premotor Cortex Exhibit “Thin Spikes”: Implications for Cell Type Classification. J. Neurosci. 31, 14235–14242 (2011).

[37] Constantinople, C. M., Disney, A. A., Maffie, J., Rudy, B. & Hawken, M. J. Quantitative analysis of neurons with Kv3 potassium channel subunits, Kv3.1b and Kv3.2, in macaque primary visual cortex. J. Comp. Neurol. 516, 291–311 (2009).

[38] Fuentealba, P. et al. Ivy Cells: A Population of Nitric-Oxide-Producing, Slow-Spiking GABAergic Neurons and Their Involvement in Hippocampal Network Activity. Neuron 57, 917–929 (2008).

